## Supplementary Figures and Legends for "A tripartite organelle platform links growth factor receptor signaling to mitochondrial metabolism"

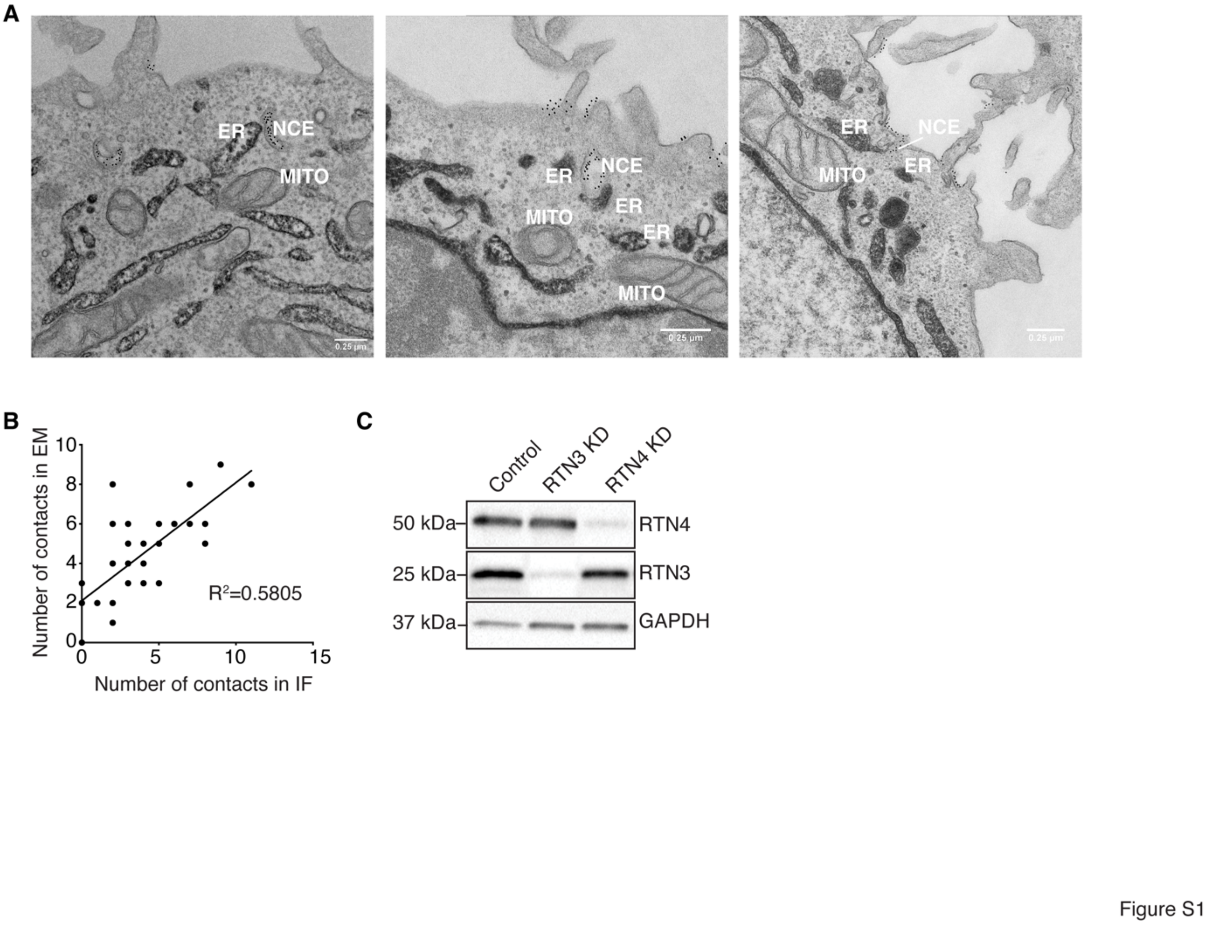


**Fig. S1. EGFR-NCE involves tripartite PM-ER-mitochondria contact sites: additional data to Figure 1. (A)** Immuno-EM micrographs showing the close spatial relationship between NCE tubules (CD147-10 nm gold), the ER (HRP-KDEL/DAB), and mitochondria in HeLa cells treated with high dose EGF for 5 mins. Bars 0.25 μM. (**B**) Correlation analysis of ER-mitochondria contacts on CLEM images. For each CLEM image pair, the number of contacts revealed by ImageJ JACoP plugin in the fluorescent image was plotted against the number of ER-mitochondria contacts (<20nm) observed in the corresponding EM image. $R^2$ = correlation coefficient. (**C**) Efficiency of RTN3 and RTN4 KD in HeLa cells was analyzed by IB. Control cells were mock treated. GAPDH, loading control. MW markers shown on the left.

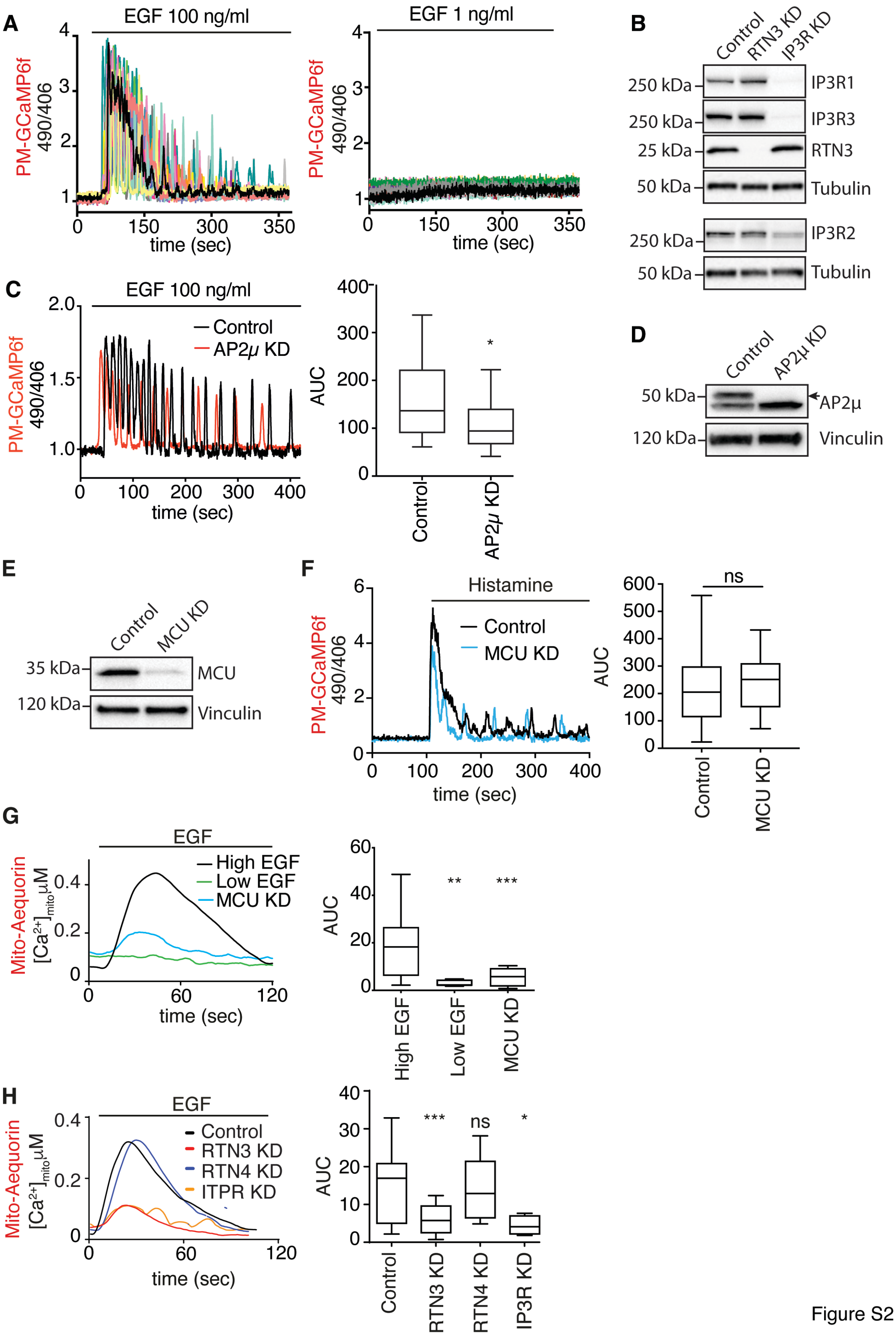
A
EGF 100 ng/ml
EGF 1 ng/ml
PM-GCaMP6f 490/406
time (sec)
B
Control
RTN3 KD
IP3R KD
250 kDa
IP3R1
250 kDa
IP3R3
25 kDa
RTN3
50 kDa
Tubulin
250 kDa
IP3R2
50 kDa
Tubulin
C
EGF 100 ng/ml
Control
AP2μ KD
AUC
D
Control
AP2μ KD
50 kDa
AP2μ
120 kDa
Vinculin
E
Control
MCU KD
35 kDa
MCU
120 kDa
Vinculin
F
Histamine
Control
MCU KD
ns
G
EGF
High EGF
Low EGF
MCU KD
Mito-Aequorin
[Ca2+]mito μM
H
EGF
Control
RTN3 KD
RTN4 KD
ITPR KD
IP3R KD

Figure S2

**Fig. S2. EGF-induced $Ca^{2+}$ response at the PM and inside mitochondria: additional data to Figure 2.** (**A**) HeLa PM-GCaMP6f cells were stimulated with low or high dose EGF. The $Ca^{2+}$ response was monitored by measuring fluorescence over time. Results are presented as the ratio of the emission at 490/406 nm for individual cells (represented by different colors). (**B**) Efficiency of RTN3 KD and IP3R KD (IP3-R1, IP3-R2, IP3-R3) in HeLa cells was analyzed by IB. Tubulin, loading control. MW markers shown on the left. (**C**) The $Ca^{2+}$ response was measured in HeLa cells subjected or not to AP2μ-KD and stimulated with high EGF as in (A). Results are presented as the ratio of the emission at 490/406 nm. Left, representative single cells traces. Right, mean AUC ± SD. Control/High EGF N=18, AP2μ KD/High EGF N=18, N=number of cells; n=1 biological replicate. (**D**) Efficiency of AP2μ KD in HeLa cells analyzed by IB (the arrow indicates the specific bands). Vinculin, loading control. MW markers shown on the left. (**E-F**) Additional controls for main Figure 2C. (**E**) Efficiency of MCU KD in HeLa cells analyzed by IB. Vinculin, loading control. MW markers shown on the left. (**E**) Histamine induces a $Ca^{2+}$ response independently of MCU. HeLa PM-GCaMP6f cells were subjected to MCU KD and stimulated with histamine (100 μM). The kinetics of the $Ca^{2+}$ response was monitored by measuring fluorescence. Results are presented as (C). Control/Histamine N=91, MCU KD/Histamine N=85, a representative experiment out of two independent replicates is shown. (**F**) Efficiency of MCU KD in HeLa cells analyzed by IB. Vinculin, loading control. MW markers shown on the left. (**G**) HeLa cells expressing Aequorin targeted to the mitochondria (Mito-Aequorin) were stimulated with low EGF or high EGF alone or with MCU KD. $Ca^{2+}$ waves inside mitochondria were detected by luminescence. Left, representative traces. Right, mean AUC ± SD. Low EGF: N=6, n=3; High EGF: N=29, n=9; High EGF/MCU KD: N=15, n=5. N=number of coverslips (whole cell population); n=biological replicates. (**H**) HeLa cells expressing Mito-Aequorin were subjected or not to the indicated KDs and stimulated with high EGF (100 ng/ml), during luminescence recording. Results are presented as in (F). Control/High EGF: N=25 n=6; RTN3 KD/High EGF: N=16, n=5; RTN4 KD/High EGF: N=11, n=5; IP3R KD/High EGF: N=4, n=2. All panels, P-value (Each Pair Student's t-test, two-tailed): *, <0.05; **, <0.01; ***, <0.001, ns, not significant.

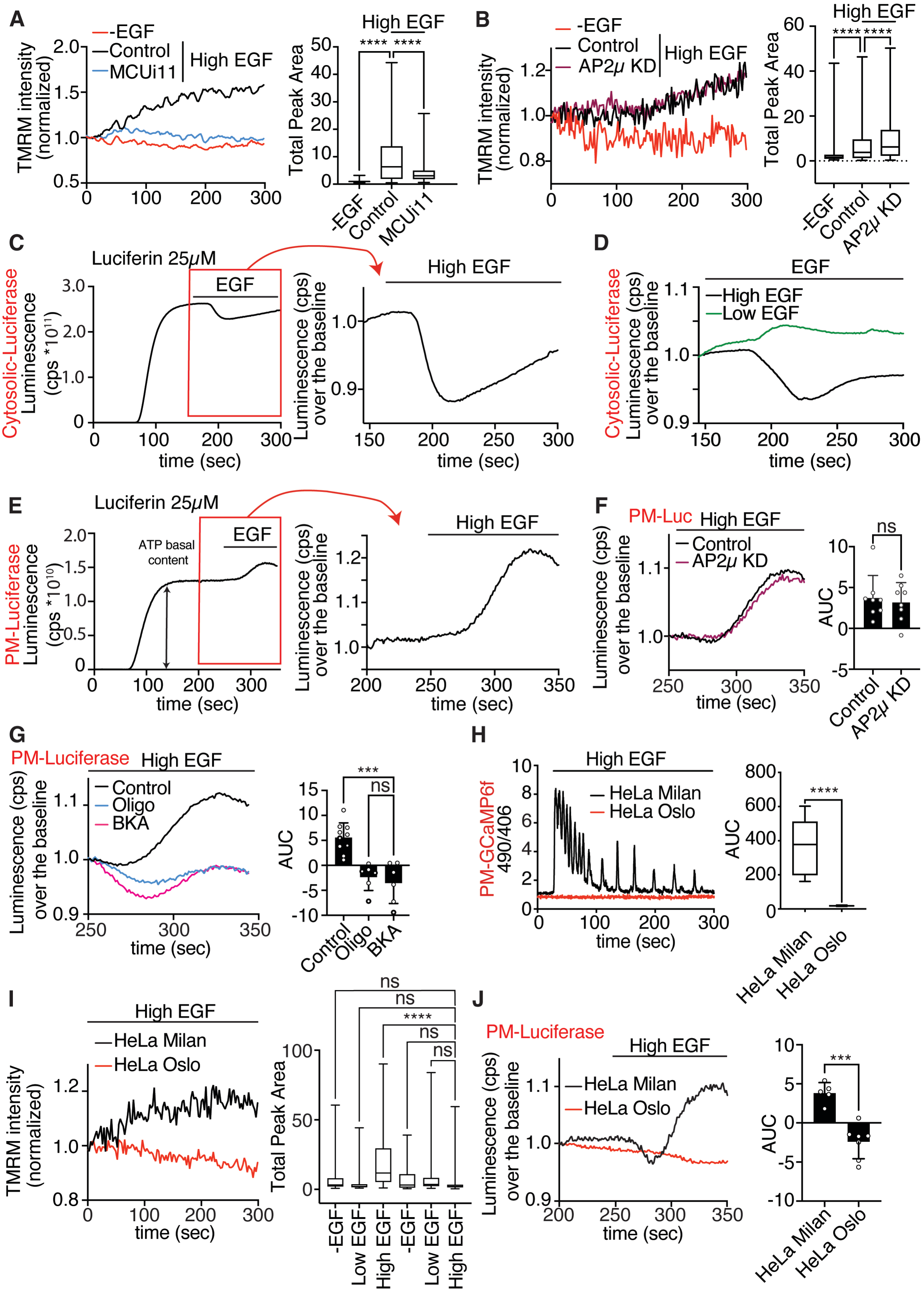
A
-EGF
Control
MCUi11
High EGF
TMRM intensity (normalized)
Total Peak Area
****
B
AP2μ KD
C
Luciferin 25μM
EGF
Cytosolic-Luciferase
Luminescence (cps *10^11)
time (sec)
Luminescence (cps) over the baseline
D
Low EGF
E
ATP basal content
PM-Luciferase
Luminescence (cps *10^10)
F
PM-Luc
ns
AUC
G
Oligo
BKA
***
H
PM-GCaMP6f 490/406
HeLa Milan
HeLa Oslo
I
J

Figure S3

**Fig. S3. Characterization and specificity of EGF-induced $Ca^{2+}$ oscillations and mitochondrial ATP production: additional data to Figure 3.** (**A**) HeLa cells were pre-treated with MCU inhibitor (MCUi11, 50 µM) or vehicle control for 30 min, then labeled with TMRM and either left unstimulated (-EGF) or stimulated high EGF (100 ng/ml). Fluorescence was recorded over 5 min. Left, fluorescence TMRM intensity. Right, mean Total peak area ± SD. -EGF N=50, Control/High EGF N=140, MCUi11/High EGF N=134, N=number of cells; n=1. (**B**) TMRM fluorescence measured in HeLa cells, subjected to AP2µ KD or mock control and stimulated high dose EGF for 5 min. Unstimulated mock cells (-EGF) served as the negative control. Left, time course of TMRM fluorescence intensity. Right, Total peak area ± SD. -EGF N=156 (n=1), Control N=823 (n=3), AP2µ KD N=825 (n=3), N=number of cells (n=3). Note that AP2 KD caused an increase in TMRM fluorescence relative to the high EGF control. (**C**) HeLa cells expressing cytosolic-luciferase were incubated with 25 µM luciferin and basal luminescence was recorded (representing an estimation of the basal ATP level in the cytosol). EGF was perfused after 250 s and reached cells after approximately 30 sec. Left, representative complete luminescence time course. Right, magnification of the red boxed area, showing luminescence levels over the baseline just before and after the addition of EGF; cps, count per seconds. (**D**) HeLa cells expressing cytosolic-luciferase were treated as in "C" and stimulated with low and high dose EGF. Median curves of luminescence over the baseline after the addition of EGF are shown. Low EGF N=8, High EGF N=19, N=number of coverslips (whole cell population). (**E**) HeLa cells expressing PM-Luc were incubated with 25 µM luciferin and basal luminescence was recorded. EGF (high dose) was perfused after 250 s and reached cells after approximately 30 sec. Left, representative complete luminescence time course. Right, magnification of the red boxed area. (**F**) Luminescence was measured in HeLa cells expressing PM-luc and subjected or not to AP2µ KD. Cells were treated with 25 µM luciferin. EGF (high dose) was perfused after 250 s and reached cells after approximately 30 sec. Left, median curves of luminescence over the baseline are shown. Right, mean AUC ± SD. Control N=8, AP2µ KD N=8, N=number of coverslips; n=2. (**G**) HeLa cells expressing PM-Luc were treated with bongkrekic acid (BKA, 50 µM) for 15 mins before addition of 25 µM luciferin and luminescence recording. EGF (high dose) was perfused after 250 s and reached cells after approximately 30 sec. Note that data reported here for Oligomycin (OMY) are the same as in Fig. 3G. Left, median curves of luminescence over the baseline. Right, mean AUC ± SD. Control N=10, BKA N=6, OMY N=6, N=number of coverslips (whole cell population); n=2, (**H-J**) HeLa Oslo cells, a clone lacking NCE but has CME, is unable to activate an EGFdependent $Ca^{2+}$ signaling at the PM and the mitochondrial response. (**H**) HeLa Milan (clone used in experiments throughout) and HeLa Oslo cells expressing PM-GCaMP6f were stimulated with high dose EGF. The kinetics of the $Ca^{2+}$ response was monitored by measuring fluorescence. Results are presented as the ratio of the emission at 490/406 nm. Left, representative single cell response curves. Right, mean AUC ± SD. Milan N=12, Oslo N=12, N=number of cells; a representative experiment of three independent biological replicates is shown. (**I**) HeLa Milan and Oslo cells labeled with TMRM were left unstimulated (-EGF) or stimulated with low or high EGF and fluorescence recorded. Left, representative time course of fluorescence intensity for cells stimulated with high EGF. Right, total peak area ± SD. HeLa Milan: -EGF N=165, Low EGF N=147, High EGF N=322. HeLa Oslo: -EGF N=39, Low EGF N=173, High EGF N=146, N=number of cells; a representative experiment of two independent biological replicates is shown. (**L**) Luminescence was recorded in HeLa Milan and Oslo cells expressing PM-Luc and treated with 25 μM luciferin. EGF (high dose) was perfused after 250 s and reached cells after approximately 30 sec. Left, median curves of luminescence over the baseline before and after the addition of EGF are shown. Right, mean AUC ± SD. HeLa Milan N=5, HeLa Oslo N=7; n=1. All panels, P-value (Each Pair Student's t-test, two-tailed): *, <0.05; **, <0.01; ***, <0.001; ****, <0.0001; ns, not significant.

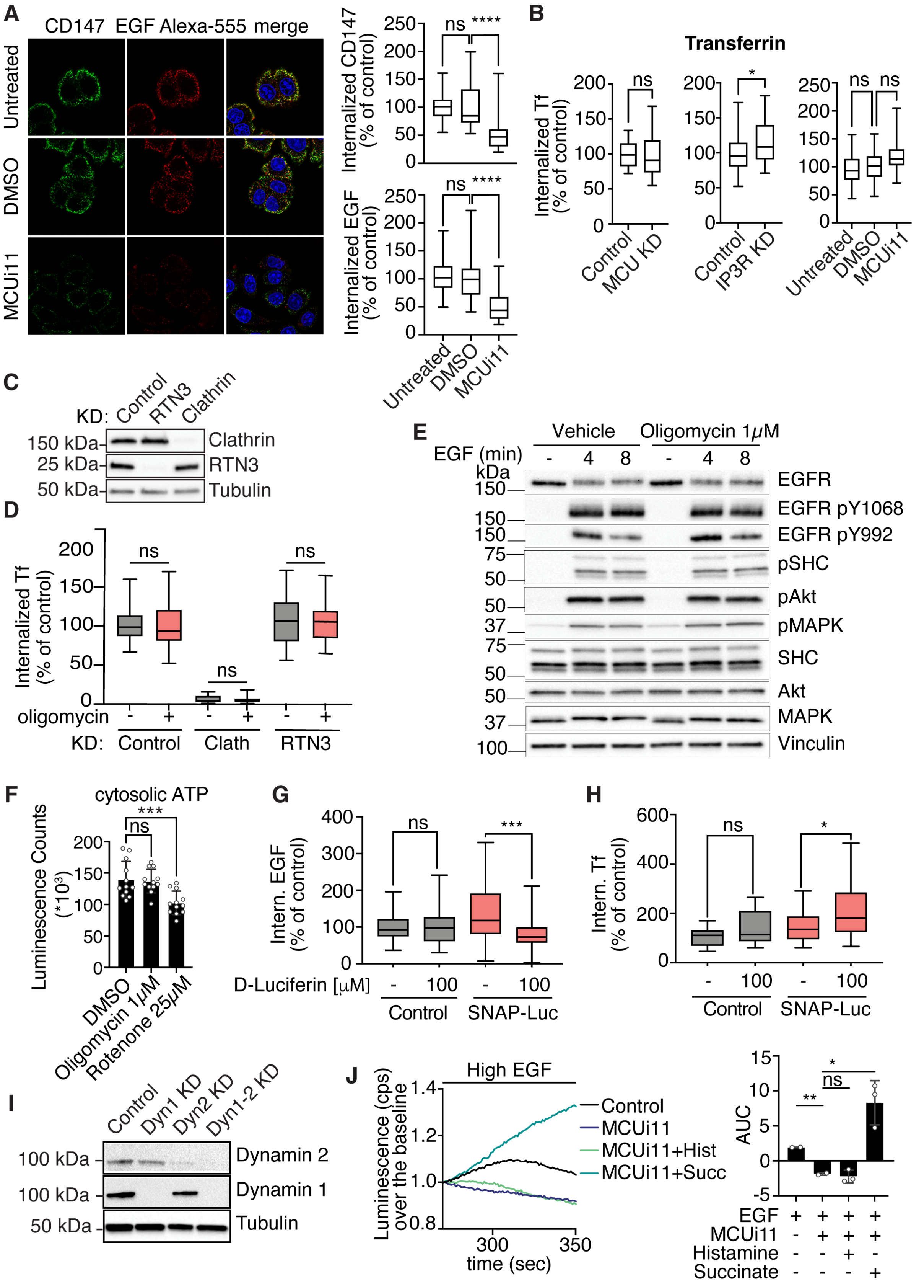
A
CD147 EGF Alexa-555 merge
Untreated
DMSO
MCUi11
Internalized CD147 (% of control)
ns ****
Internalized EGF (% of control)
ns ****
Untreated
DMSO
MCUi11
B
Transferrin
Internalized Tf (% of control)
ns
Control
MCU KD
*
Control
IP3R KD
ns ns
Untreated
DMSO
MCUi11
C
KD: Control RTN3 Clathrin
150 kDa Clathrin
25 kDa RTN3
50 kDa Tubulin
D
Internalized Tf (% of control)
ns
ns
ns
oligomycin - + - + - +
KD: Control Clath RTN3
E
Vehicle
Oligomycin 1μM
EGF (min) - 4 8 - 4 8
kDa
EGFR
EGFR pY1068
EGFR pY992
pSHC
pAkt
pMAPK
SHC
Akt
MAPK
Vinculin
F
cytosolic ATP
Luminescence Counts (*10³)
***
ns
DMSO
Oligomycin 1μM
Rotenone 25μM
G
Intern. EGF (% of control)
ns
***
D-Luciferin [μM] - 100 - 100
Control SNAP-Luc
H
Intern. Tf (% of control)
ns
*
- 100 - 100
Control SNAP-Luc
I
Control Dyn1 KD Dyn2 KD Dyn1-2 KD
100 kDa Dynamin 2
100 kDa Dynamin 1
50 kDa Tubulin
J
High EGF
Luminescence (cps) over the baseline
time (sec)
Control
MCUi11
MCUi11+Hist
MCUi11+Succ
AUC
**
ns
*
EGF + + + +
MCUi11 - + + +
Histamine - - + -
Succinate - - - +

Figure S4

**Fig. S4. EGF-induced $Ca^{2+}$ waves and mitochondrial ATP production are required for NCE execution: additional data to Figure 4.** (**A**) HeLa cells treated with MCUi11 (50 µM for 30 min), DMSO vehicle control or left untreated, were incubated *in vivo* with a specific anti-CD147 antibody for 60 min at 4°C, then with an Alexa-488 secondary antibody (green) for 30 min at 4°C and then stimulated with high dose Alexa647-EGF (100 ng/ml, red) for 5 min at 37°C. Cells were subjected to acid wash treatment prior to fixation to visualize only internalized CD147/EGF. MCUi11 was kept during stimulation. Blue, DAPI. Bar, 10 µm. Right, Quantification of internalized CD147 (top) and EGF (bottom). Mean integrated fluorescence intensity ± SD is reported as a percentage relative to control. N=number of cells: untreated N=116, DMSO N=110, MCUi11 N=115 (n=2). (**B**) Transferrin (Tf) internalization in HeLa cells subjected to the indicated KDs and treatments. Alexa647-Tf internalization was followed for 8 min at 37°C in HeLa cells. Mean integrated fluorescence intensity ± SD is reported as a percentage relative to control. N=number of cells: Control N=113, MCU KD N=98 (n=2); Control N=396, IP3R KD N=324 (n=4); untreated N=134, DMSO N=135, MCUi11 N=137 (n=2). (**C**) Efficiency of RTN3 and clathrin KD in HeLa cells, analyzed by IB. Tubulin, loading control. MW markers shown on the left. (**D**) Transferrin (Tf) internalization in HeLa cells subjected to OMY treatment or RTN3 KD or clathrin KD was in measured and reported as in (A). N=number of cells: Control N=244, Control+OMY=250 (n=4); Clathrin KD N=231, Clathrin KD+OMY N=209, (n=4); RTN3 KD N=185, RTN3 KD+OMY N=183 (n=3). (**E**) IB analysis of the expression and phosphorylation status of EGFR and the indicated signaling effectors in HeLa cells, treated with OMY (1 µM, pretreatment for 5 min, as in main Figs. 4B, D) or vehicle and stimulated or not with high EGF dose (100 ng/ml) for different timepoints. Vinculin, loading control. MW markers shown on the left. Note that acute OMY treatment does not affect EGFR activation nor its downstream signaling, suggesting that the inhibition of NCE is independent of EGFR activation. (**F**) Measurement of the total cellular ATP content (measured with ADP/ATP Ratio Assay Kit, ab65313, abcam) upon acute treatment with OMY (1 µM, pretreatment for 5 min). Chronic treatment with rotenone (50 µM for 4 h) was used as positive control of cytosolic ATP reduction. Note that, acute OMY treatment does not affect the total ATP content while affecting EGF/CD147 NCE (**Fig. 4B**). For each condition, n=4. (**G, H**) HeLa cells were transfected with PM-Luc and treated or not with luciferin (100 µM). Internalization of EGF (G) or Tf (H) was monitored for 5 min at 37°C. Relative EGF/Tf fluorescence intensity is expressed as % of control ± SD. N=number of cells for EGF: Control N=158 (n=3); Control+Luciferin N=142 (n=3); SNAP-

Luc N=116 (n=4); SNAP-Luc+Luciferin N=78 (n=4). N=number of cells for Tf: Control N=294 (n=4); Control+Luciferin N=270 (n=4); SNAP-Luc N=129 (n=4); SNAP-Luc+Luciferin N=136 (n=4). (**I**) Efficiency of Dynamin 1 and Dynamin II KDs in HeLa cells (used in Fig. 4D, E), as compared to mock-treated control, analyzed by IB. Tubulin, loading control. MW markers shown on the left. (**J**) Additional controls for main Fig. 4F. Succinate treatment rescues the PM-localized ATP increase induced by high EGF in MCU inhibited cells. HeLa cells expressing PM-Luc were left untreated or treated with MCUi11 (50 μM for 30 mins and kept during the recording) alone or in combination with histamine (100 μM) or succinate (5 mM), and then were stimulated with high EGF (100 ng/ml), during luminescence recording. Left, representative curves of luminescence over the baseline after the addition of EGF; cps, count per second. Right, mean AUC ± SD. N=number of coverslips (whole cell population): Control N=2, MCUi11 N=2, MCUi11/Histamine N=3, MCUi11/Succinate N=3 (n=1). All panels, P-value (Each Pair Student's t-test, two-tailed): *, <0.05; **, <0.01; ***, <0.001; ****, <0.0001; ns, not significant.

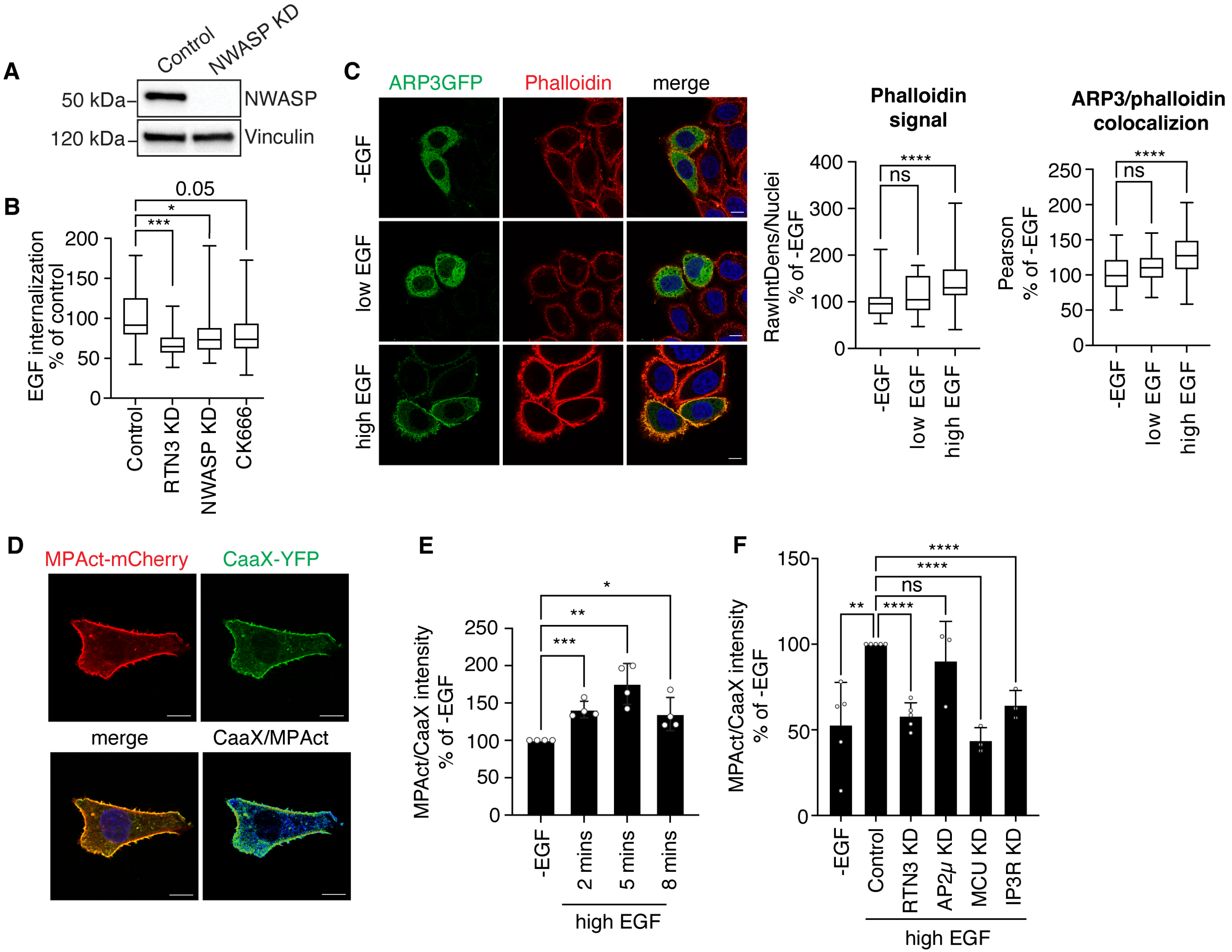
A
Control
NWASP KD
50 kDa
120 kDa
NWASP
Vinculin
B
EGF internalization
% of control
0.05
*
***
Control
RTN3 KD
NWASP KD
CK666
C
ARP3GFP
Phalloidin
merge
-EGF
low EGF
high EGF
Phalloidin signal
RawIntDens/Nuclei
% of -EGF
****
ns
ARP3/phalloidin colocalizion
Pearson
% of -EGF
D
MPAct-mCherry
CaaX-YFP
merge
CaaX/MPAct
E
MPAct/CaaX intensity
% of -EGF
-EGF
2 mins
5 mins
8 mins
high EGF
F
MPAct/CaaX intensity
% of -EGF
-EGF
Control
RTN3 KD
AP2μ KD
MCU KD
IP3R KD
high EGF

Figure S5

**Fig. S5. Regulation of F-actin and ARP3 localization by EGF, and characterization of the cortical actin probe, MPAct: additional data to Figure 5.** (**A**) Efficiency of NWASP KD in HeLa cells analyzed by IB. GAPDH, loading control. MW markers shown on the left. (**B**) EGF internalization in HeLa cells subjected to the indicated KDs and high dose EGF stimulation (same experiment as in main Figure 5A). Alexa647-EGF internalization was followed for 5 min at 37°C. Cells were subjected to acid wash treatment prior to fixation to visualize only internalized EGF. Mean integrated fluorescence intensity ± SD is reported as a percentage relative to control. RTN3 KD was used as a positive control for NCE inhibition. N=number of cells: Control N=258, RTN3 KD N=73, NWASP KD N=231, CK666 N=242 (n=3). (**C**) HeLa cells transfected with GFP-ARP3 (green) were stimulated with low or high dose of EGF or left untreated. After fixation, cells were stained with phalloidin-TRITC (red). Left, representative IF images, Blue, DAPI. Bar, 10 μm. Middle, raw integrated density of the phalloidin signal/nuclei ± SD is reported as a percentage relative to the -EGF sample. Right, Pearson colocalization coefficient between phalloidin and ARP3 reported as a percentage relative to the -EGF sample. N=number of cells: -EGF N=200, Low EGF N=189, High EGF N=238 (n=3). (**D**) HeLa cells were subjected to MPAct-mCherry and YFP-CaaX co-transfection. A representative IF image split into the different channels, as indicated, is shown. Blue, DAPI. Bar, 10 μm. (**E**) Ratiometric analysis of MPA density in HeLa cells subjected to MPAct-mCherry and YFP-CaaX co-transfection and then stimulated with high EGF dose (100 ng/ml) or left unstimulated for the indicated time points. Mean raw integrated fluorescence intensity ± SD is reported as a percentage relative to -EGF cells. N=number of cells: -EGF N=164, High EGF/2 mins N=220, High EGF/5 mins N=178, High EGF/8 mins N=157 (n=4). (**F**) Ratiometric analysis of MPA density in HeLa cells subjected to the indicated KDs, followed by MPAct-mCherry and YFP-CaaX co-transfection and then stimulated with high EGF dose (100 ng/ml). Mean raw integrated fluorescence intensity ± SD is reported as a percentage relative to -EGF cells. N=number of cells: -EGF N=220, Control/High EGF N=239, RTN3 KD N=249, AP2μ KD N=188, MCU KD N=218, IP3R KD N=196, n=3. All panels, P-value (Each Pair Student's t-test, two-tailed): *, <0.05; **, <0.01; ***, <0.001; ****, <0.0001; ns, not significant.

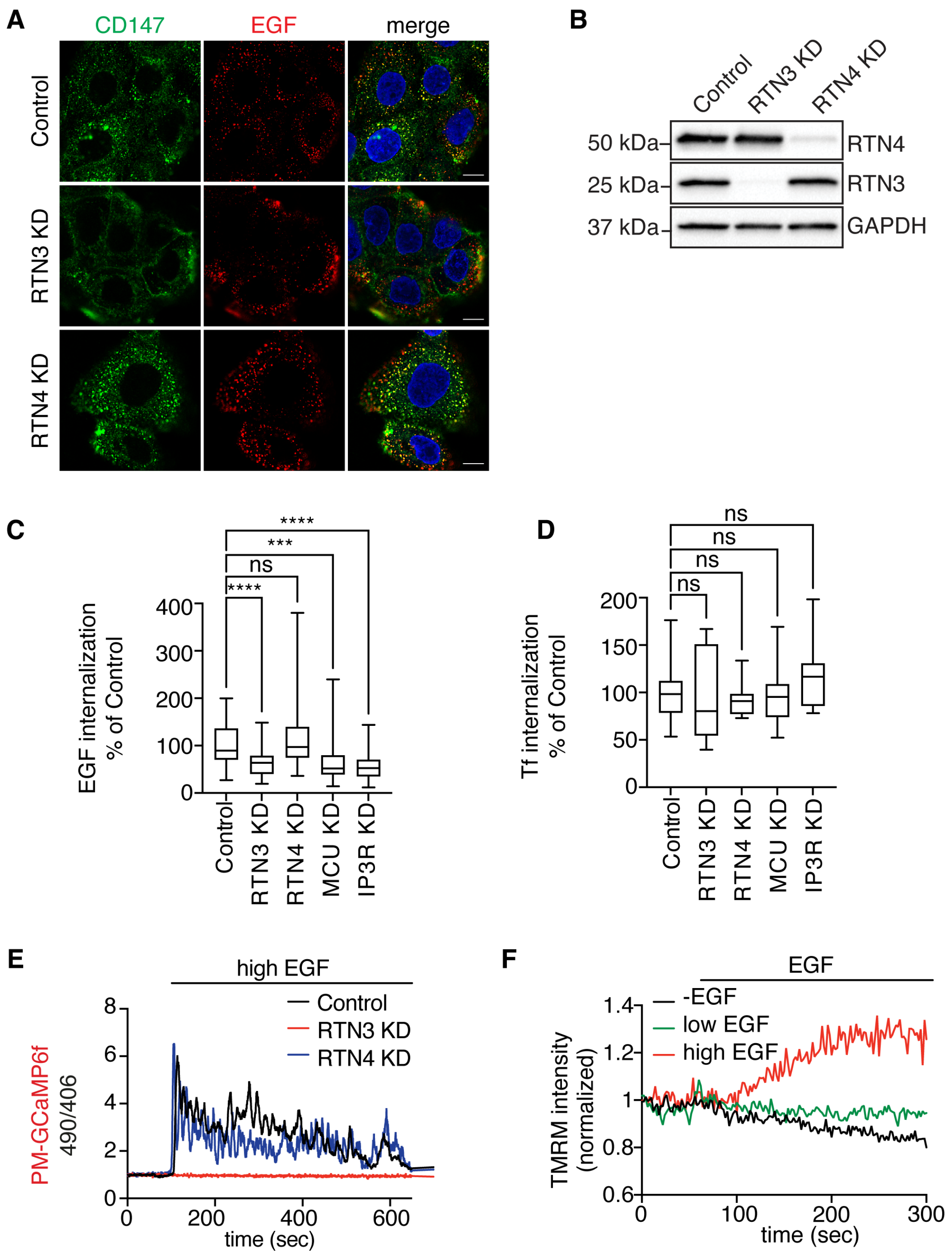
A
CD147
EGF
merge
Control
RTN3 KD
RTN4 KD
B
Control
RTN3 KD
RTN4 KD
50 kDa
25 kDa
37 kDa
RTN4
RTN3
GAPDH
C
EGF internalization
% of Control
ns
Control
RTN3 KD
RTN4 KD
MCU KD
IP3R KD
D
Tf internalization
% of Control
E
high EGF
PM-GCaMP6f
490/406
Control
time (sec)
F
EGF
-EGF
low EGF
high EGF
TMRM intensity
(normalized)

Figure S6

**Fig. S6. Characterization of EGFR-NCE in the HaCaT cell model system.** (**A**) CD147 internalization was monitored *in vivo* by IF in HaCaT cells subjected to the indicated KDs or mock transfection and stimulated with high dose EGF. Cells were incubated with an anti-CD147 antibody for 60 min at 4°C, then with an Alexa-488 secondary antibody (green) for 30 min at 4°C before the addition of high dose Alexa647-EGF (red) for 12 min at 37°C. Cells were subjected to acid wash treatment prior to fixation to remove membrane-bound antibodies and analyzed by confocal microscopy. Blue, DAPI. Bar, 10 μm. Quantification is reported in main Fig. 6A. (**B**) Efficiency of RTN3 and RTN4 KD in HaCaT cells analyzed by IB. GAPDH, loading control. MW markers shown on the left. (**C**) EGF internalization was monitored in HaCaT cells subjected to the indicated KDs or mock transfection (Control) and stimulated with high dose Alexa647-EGF. Quantification of relative EGF fluorescence intensity is expressed as % of control ± SD. N=number of cells: Control N=301 (n=6), RTN3 KD N=97 (n=3), RTN4 KD N=179 (n=5), MCU KD N=253 (n=5), IP3R KD N=199 (n=4). P-value (Each Pair Student's t-test two-tailed): ***, <0.001, ****, <0.0001; ns, not significant. (**D**) Alexa647-Tf internalization was followed for 12 min at 37°C in HaCaT cells subjected to the indicated KDs or mock transfection (Control). Mean integrated fluorescence intensity ± SD is reported as a percentage relative to control. N=number of cells: Control N=54, RTN3 KD N=44, RTN4 KD N=45, MCU KD N=58, IP3R KD N=47 (n=1). ns, not significant. (**E**) HaCaT cells stably expressing PM-GCaMP6f were stimulated with high dose EGF in the presence of transient RTN3 or RTN4 KD or mock control. The $Ca^{2+}$ response was monitored by measuring fluorescence and results are presented as the ratio of the emission at 490/406 nm. Representative single cell curves are reported. Quantitation is shown in main Fig. 6C. (**F**) HaCaT cells were labeled with TMRM and left unstimulated (- EGF) or stimulated with low or high EGF. Fluorescence TMRM intensity is reported. Quantitation is shown in main Fig. 6D.

**Movies**

**Movie S1 (supplementary to Fig. 1B). Tomographic 3D reconstruction showing a tripartite PM-ER-mitochondria contact site.** The cortical region of a HeLa cell treated with high dose EGF for 5 min and analyzed by immuno-EM as in Figure 1A is shown. The spatial proximity between internalizing NCE structures (light grey) containing the cargo CD147 (yellow dots), the ER (blue) and mitochondria (MITO, green) is visible. The ER was automatically recognized through HRP-KDEL staining, while mitochondria were manually segmented.

**Movie S2 (supplementary to Fig. 1B).** ER tubules interposed between mitochondria and CD147-positive TI. Single-tilted tomographic reconstruction (see Materials and Methods) of ER tubules in contact with TIs—internalizing gold-labeled CD147—and mitochondria. Note the contact of ER (pseudocolored in blue) profiles recognized through HRP-KDEL staining with mitochondria (pseudocolor in green) and with CD147-internalizing TI (labeled in gold). Single-tilted tomographic reconstruction, shown in Fig. 1C, was extracted from this movie.

**Movie S3 (supplementary to Fig. 2B). EGF-induced $Ca^{2+}$ oscillations at the PM in HeLa cells detected by PM-GCaMP6f.** Cells treated with low EGF (left), High EGF (centre), High EGF + RTN3 KD (right) (5 frame/sec). EGF was added after 45 seconds from the start of the recording. The movie was cut at 300 seconds and accelerated at 75 frame/sec (15x).

**Movie S4 (supplementary to Fig. 2E). EGF-induced $Ca^{2+}$ oscillations inside the mitochondria in HeLa cells detected by mito-GCaMP6m.** Cells treated with low EGF (left) or high EGF (right) (5 frame/sec). EGF was added after 45 seconds from the start of the recording. The movie was cut at 300 seconds and accelerated at 75 frame/sec (15x).

**Movie S5 (supplementary to Fig. 3A). EGF-induced increase in $\Delta\Psi m$.** Change in TMRM fluorescence observed in unstimulated HeLa cells (left), or cells stimulated with low (center) or high (right) dose EGF (0.5 frame/sec). EGF was added after 45 seconds from the start of the recording. The movie was accelerated at 50 frame/sec (100x).

**Movie S6 (supplementary to Fig. 6E). High dose EGF promotes collective cell migration in a wound-healing assay.** Time-lapse video microscopy showing wound healing in sub-confluent HaCaT cells, unstimulated (left), or treated with low (middle) or high (right) dose EGF for 30 h (1 frame/5 min). The movie was accelerated at 50 frame/sec (15000x).
